# Discovery of large genomic inversions using pooled clone sequencing

**DOI:** 10.1101/015156

**Authors:** Marzieh Eslami Rasekh, Giorgia Chiatante, Mattia Miroballo, Joyce Tang, Mario Ventura, Chris T. Amemiya, Evan E. Eichler, Francesca Antonacci, Can Alkan

## Abstract

**Motivation:** There are many different forms of genomic structural variation that can be broadly classified into two groups as copy number variation (CNV) and balanced rearrangements. Although many algorithms are now available in the literature that aim to characterize CNVs, discovery of balanced rearrangements (inversions and translocations) remains an open problem. This is mainly because the breakpoints of such events typically lie within segmental duplications and common repeats, which reduce the mappability of short reads. The 1000 Genomes Project spearheaded the development of several methods to identify inversions, however, they are limited to relatively short inversions, and there are currently no available algorithms to discover large inversions using high throughput sequencing technologies (HTS).

**Results:** Here we propose to use a sequencing method (Kitzman *et al.*, 2011) originally developed to improve haplotype phasing to characterize large genomic inversions. This method, called pooled clone sequencing, merges the advantages of clone based sequencing approaches with the speed and cost efficiency of HTS technologies. Using data generated with pooled clone sequencing method, we developed a novel algorithm, dipSeq, to discover large inversions (>500 Kbp). We show the power of dipSeq first on simulated data, and then apply it to the genome of a HapMap individual (NA12878). We were able to accurately discover all previously known and experimentally validated large inversions in the same genome. We also identified a novel inversion, and confirmed using fluorescent in situ hybridization.

**Availability:** Implementation of the dipSeq algorithm is available at https://github.com/BilkentCompGen/dipseq

## 1 INTRODUCTION

Genomic structural variants are defined as alterations in the DNA that affect >50 bp that may delete, insert, duplicate, invert, or move genomic sequence (Alkan *et al.*, 2011). Structural variation (SV) is shown to be common in human genomes (Iafrate *et al.*, 2004; Sebat *et al.*, 2004), which caused increased interest in the characterization of both normal (Tuzun *et al.*, 2005; Kidd *et al.*, 2008; Mills *et al.*, 2011), and disease-causing large variants (Sharp *et al.*, 2006; Antonacci *et al.*, 2010). Furthermore, SVs are known to be one of the driving forces of creation of new haplotypes (Steinberg *et al.*, 2012), and evolution (Ventura *et al.*, 2011).

Copy number variations (CNVs) were initially identified using BAC (bacterial artificial chromosome) and oligo array comparative genomic hybridization (CGH) (Iafrate *et al.*, 2004; Sebat *et al.*, 2004; Redon *et al.*, 2006; Conrad *et al.*, 2010), and SNP genotyping arrays (Redon *et al.*, 2006; McCarroll *et al.*, 2006). A more detailed map of SV was made possible using fosmid end sequencing (Tuzun *et al.*, 2005; Kidd *et al.*, 2008), however this method was too expensive and time-consuming since it involved creating and plating of fosmid libraries followed with Sanger sequencing. Introduction of high throughput sequencing (HTS) finally made it possible to screen the genomes of many (Korbel *et al.*, 2007; Alkan *et al.*, 2009; Hormozdiari *et al.*, 2009; Yoon *et al.*, 2009) to thousands (Mills *et al.*, 2011) of individuals.

Although there are now many algorithms to discover and genotype SV using HTS data (Medvedev *et al.*, 2009; Alkan *et al.*, 2011), they mainly focus on CNVs, which change the amount of DNA, such as deletions, duplications, insertions, and transpositions. Other types of SV, namely *balanced rearrangements* such as inversions and translocations are harder to detect due to the fact that their breakpoints usually lie within complex repeats, reducing mappability. Balanced rearrangements also do not alter the read depth, which makes the use of read depth signature (Medvedev *et al.*, 2009; Yoon *et al.*, 2009; Alkan *et al.*, 2009) irrelevant for their detection. Therefore, very few attempts to characterize inversions are reliable only for small inversions (~10-50 Kbp) (Hormozdiari *et al.*, 2009; Rausch *et al.*, 2012; Layer *et al.*, 2014; Chaisson *et al.*, 2015), and exhibit high false discovery rates in translocation call sets (Talkowski *et al.*, 2011). Another algorithm, GASVPro (Sindi *et al.*, 2012) is also able to detect inversions with a size limit up to 500 Kbp, however its sensitivity and specificity for large inversions are yet untested. Characterization of larger genomic inversions using HTS remains an open problem.

## 2 BACKGROUND AND MOTIVATION

Most known examples of large inversions have come indirectly from studies on human disease where inversions have no detectable effect in parents, but increase the risk of a disease-associated rearrangement in the offspring. In the Williams-Beuren syndrome, for example, 25-30% of transmitting parents have a 1.5 Mbp inversion encompassing the commonly deleted region, whereas the same inversion is present in only 6% of the general population (Osborne *et al.*, 2001). Similarly, a polymorphic inversion has been reported at 15q11-q13 and is seen more frequently in mothers who transmit de novo deletions resulting in the Angelman syndrome (Gimelli *et al.*, 2003). Two more striking examples are found in the Sotos syndrome (Visser *et al.*, 2005) and the 17q21.31 microdeletion syndrome (Stefansson *et al.*, 2005; Sharp *et al.*, 2006; Koolen *et al.*, 2006; Zody *et al.*, 2008; Steinberg *et al.*, 2012). In each of these disorders, every parent studied to date in which a *de novo* microdeletion arises carries an inversion of the same region. All these inversions are enriched in segmental duplications at their breakpoints, leading to an increased susceptibility to non-allelic homologous recombination (NAHR) and risk for disease-causing rearrangements to occur in the offspring. The typical presence of duplicated sequences at the inversion boundaries is also the major challenge for inversion detection.

The development of a map of inversion polymorphisms will provide valuable information regarding their distribution and frequency in the human genome and will be important for future studies aimed to unravel how inversions and the segmental duplications architecture associated with inverted haplotypes contribute to genomic susceptibility to disease rearrangements.

The common method to discover inversions is to analyze the read pair signature (Medvedev *et al.*, 2009; Alkan *et al.*, 2011), where the mapping strand of the read pairs spanning the inversion breakpoints will be different from what is expected (Figure 1). For example, the Illumina platform generates read pairs from opposing strands, however, if the DNA fragment spans an inversion breakpoint, they will both be mapped to the same strand. They will also be separated from each other by a distance approximately same with the inversion size. When the inversion is large, the *real* mapping distance between pairs also increases, therefore increasing the chance of incorrect mapping due to the common repeats that lie in between.

**Fig. 1.**
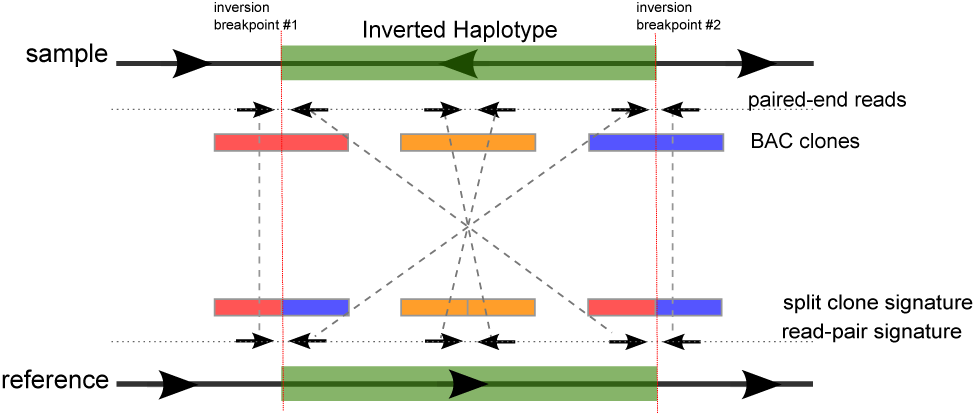
Sequence signatures used by the dipSeq algorithm. In the presence of an inverted haplotype in the sequenced genome, we look for both read pair and split clone signatures. Paired-end reads that span the inversion breakpoints will be mapped to the same strand with a large distance between them, instead of the concordant read pairs that map to opposing strands (Medvedev *et al.*, 2009; Alkan *et al.*, 2011). Large insert clones will show mapping properties similar to the split read sequence signature (Ye *et al.*, 2009), but since we do not have the full clone sequence, or sufficient coverage to assemble clones, we interrogate lengths of contiguous read mapping (Methods).

The HTS platforms generate data at very high rates with minimal cost. However, since both the HTS reads (100-150 bp for Illumina), and the DNA fragments are very short (350-500 bp), the mappability of the HTS data is dramatically reduced in repeat-rich regions that harbor most of the inversion breakpoints. On the contrary, the now-largely-abandoned method of clone-by-clone sequencing (International Human Genome Sequencing Consortium, 2001) enables data observation from much larger genomic intervals (40-to-200 Kbp), but the associated costs are substantially higher. A sequencing method, called *pooled clone sequencing* (PCS) aims to combine the advantages of clone-by-clone sequencing, with the cost and time efficiency offered by the HTS platforms (Kitzman *et al.*, 2011) (see Methods). Although pooled clone sequencing was developed to improve haplotype phasing and to characterize large haplotype blocks, we propose a novel algorithm, *dipSeq*, that utilizes PCS to discover large genomic inversions (>500 Kbp).

## 3 OBSERVATION AND APPROACH

**Pooled clone sequencing.** Kitzman *et al.* (2011) developed the pooled clone sequencing (PCS) method to improve haplotype phasing. Basically, genomic DNA is cloned into cloning vectors (fosmids, or BACs), which are then diluted to approximately 3% coverage of the diploid genome, and randomly placed into several number of pools (Methods). Next, the pools are barcoded, and sequenced using the Illumina platform. Note that, due to dilution and random generation of pools, it is expected that pools will not harbor overlapping clones within themselves (Kitzman *et al.*, 2011). We provide a method to approximately calculate the probability of having overlapping clones within a pool in the Supplementary Note.

Our approach to discover large (>500 Kbp) genomic inversion using PCS follows from the observation that, clones (BAC or fosmid) that span the inversion breakpoint will be split into two sections when mapped to the reference genome, also separated by a distance approximately the size of the inversion. We call this sequence signature as *split clones* (Figure 1, which is similar to the split read sequence signature used by several SV discovery tools such as DELLY (Rausch *et al.*, 2012) and Pindel (Ye *et al.*, 2009). Based on these observations, we developed a novel combinatorial algorithm and statistical heuristics called *dipSeq* (**d**iscover **i**nversions using **p**ooled **Seq**uencing). Briefly, dipSeq searches for both read pair and split clone sequence signatures using the mapping locations of pooled clone sequencing reads, and requires split clones from different pools to cluster at the same putative inversion breakpoints (Methods). Ambiguity due to multiple possible pairings of split clones are resolved using an approximation algorithm for the maximal quasi clique problem (Brunato *et al.*, 2008), and paired-end read support further assigns confidence score for the predicted inversion calls.

dipSeq proves its potential when tested on simulated data, and it is able to discover previously characterized large inversions (>500 Kbp) in the genome of a human individual (NA12878), using pooled BAC sequence data. dipSeq is *theoretically* compatible with all similarly constructed pooled sequence data, such as the TruSeq Synthetic Long-Reads (Moleculo) (Kuleshov *et al.*, 2014), or the Complete Genomics LFR Technology (Peters *et al.*, 2012), provided that the pooled large DNA fragment sizes follow a Gaussian distribution. However, it should be noted that, large clone size is required to span segmental duplication blocks, and smaller clones such as fosmids may not be sufficient to detect inversions around segmental duplications (Kitzman *et al.*, 2011). Therefore, the theoretical minimum inversion size detectable by dipSeq is limited by clone length, i.e. 150 Kbp when BACs are used.

## 4 METHODS

### 4.1 Building pooled clone libraries

We first generate a single whole-genome BAC library with long inserts (~140 Kbp). This procedure is a modification of the original haplotyping method previously described by Kitzman et al. (2011), that generates fosmid libraries with ~40 Kbp inserts. Here we use BAC clones, since long inserts are required to span the large duplication blocks where inversion breakpoints typically map (Kidd *et al.*, 2008; Kitzman *et al.*, 2011). We then randomly partition the library into pools such that each pool is essentially a haploid mixture of clones derived from either the maternal or paternal DNA at each genomic location. High-throughput sequencing of each pool provides haplotype information for each clone in that pool.

We used genomic DNA from a HapMap Project individual (NA12878) to construct the BAC library. High molecular weight DNA was isolated, partially EcoRI digested, and subcloned into pCC1BAC vector (Epicentre) to create a ~140 Kbp insert library using previously described protocols (Smith *et al.*, 2010). We then split a portion of this library to 3 sets of 96 pools each, with 230 clones per pool for set 1, 389 clones per pool for set 2 and 153 clones per pool for set 3. Each pool was expanded by direct liquid outgrowth after infection. We next construct 96 barcoded sequencing libraries per each set, for a total of 288 sequencing libraries (Adey *et al.*, 2010). Libraries from each set were indexed with barcodes, combined and sequenced using the Illumina HiSeq platform (101 bp paired-end reads). Upon sequencing a total of 74,112 clones (22,080 in Set 1, 37,344 in Set 2 and 14,688 in Set 3) we obtained 3.38X expected physical depth of coverage. After read mapping and clone reconstruction (Section 4.3), 87.58% of the genome was covered by one or more clones.

### 4.2 Read mapping

We first map the paired-end reads generated for each pool separately to the human reference genome assembly (GRCh37). Our dipSeq algorithm does not depend on any specific aligner, but in this study we used both BWA (Li and Durbin, 2009), and mrFAST (Xin *et al.*, 2013). We then separate the read pairs that map in the same orientation (i.e. read pair signature for inversions using Illumina), and those that map concordantly (within 4 standard deviations of the average fragment span size) into separate files to facilitate clone reconstruction (Section 4.3), and read pair support calculation (Section 4.4).

### 4.3 Reconstructing clones

We use only the concordantly mapped read pairs to infer the locations of clones. However, due to the low depth and breadth of coverage, it is not always possible to observe a continuous mapping of read pairs that collectively span genomic intervals within expected size of BAC clones. To overcome this issue, we apply several heuristics to identify clone locations. Scanning from the beginning to end of each chromosome’s reads, we first identify windows of 2 times the maximum fragment size that are covered by at least 50%. We use such regions as seeds and then extend these seed windows using any read pairs that map to its flanking regions with a distance of at most 1.5 Kbp. Although the parameters we used here may seem arbitrary, in fact they were obtained by applying an optimization grid on simulated BAC data (Section 5.1, and Supplementary Note). This algorithm runs in *O*(*n* log *n*) time for sorting the reads, and amortized run time of *O*(*n*) for reconstructing the clones, where *n* is the number of reads.

### 4.4 Inversion Discovery

After the identification of read pairs with inversion signature (i.e. mapping to same strand), and the predicted clone locations, we then look for potential split clones in each pool by pairing clones that the summation of their lengths is within an expected size range (*μ*_clone_ ± 3*σ*_clone_, where *μ*_clone_ is the mean clone size and *σ*_clone_ is the standard deviation).

We also require the distance between the split clones to be within the inversion size limits we are trying to discover. In this study we set this parameter to 500 Kbp–10 Mbp (Figure 2). Therefore, two regions r_k_ and r_l_ are predicted to be a split clone, denoted as 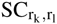 if:

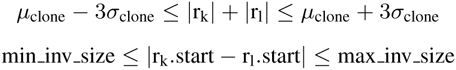

**Fig. 2.**
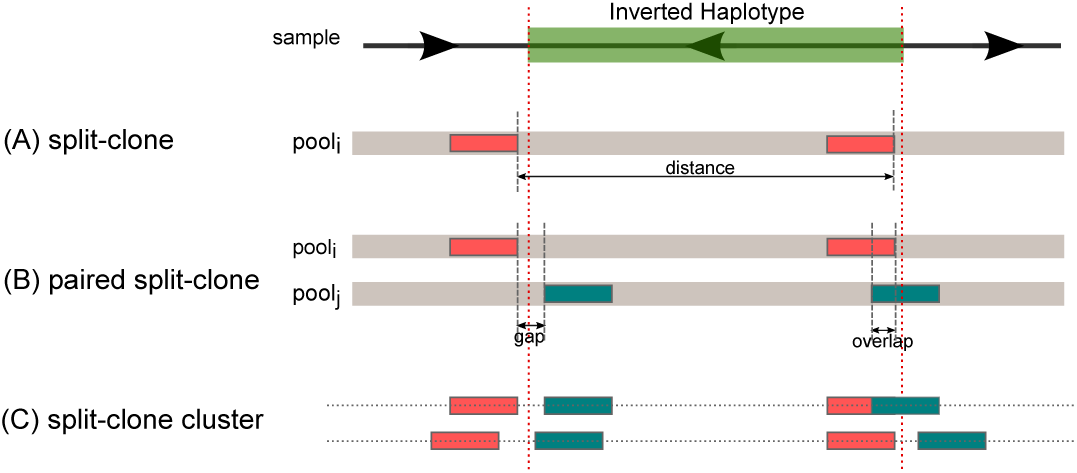
Clustering split clones to detect inversions. (A) We first identify clone locations that are shorter than the expected clone size, but when paired with another short clone found in the same pool, the total length sums up to a full clone length. We refer to such clones as “split clones”. (B) We then cluster pairs of split clones that are mapped to approximately the same breakpoints. Note that due to read mapping errors and our clone reconstruction heuristics, a split clone may be identified as spanning a breakpoint. (C) Finally we cluster multiples of split clones from different pools if they agree on breakpoint location and the size of the inversion. *gap:* size of the region between the start and end locations of split clones from different pools. *overlap:* size of the overlapping region of split clones from different pools.

Assuming the inferred clone locations are sorted by mapping locations, our algorithm can detect split clones in *O*(*n*) amortized run time, where *n* is the number of inferred clones. However, the constant coefficient increases with the increase of average sequence coverage.

We build inversion clusters by identifying two split clone pairs from different pools that are compatible (i.e. same breakpoint locations and inversion size). We denote such compatible pairs as a *pair of split clones* (PSC). Due to both mapping errors and biases caused by our sliding window approach we permit a gap or overlap between the split clones to be paired (Figure 2b). We expect the inversion breakpoints to lie between these gaps. Two split clones 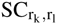 and 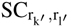, are *compatible* to be in the same paired split clone (PSC) set, assuming r_k_/r_k′_, are located upstream of r_1_/r_1′_, if:

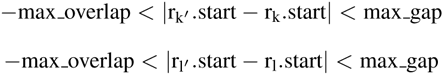

Here we set the max_gap = −1 × max_overlap = *μ*_clone_. Note that adding more split clones to the same cluster will narrow down the gap size in breakpoint estimate. However, not all of the split clones we identify signal an inversion event. In an ideal case, where there are no mapping errors, other forms of structural variation, or areas with low mappability may also show themselves as split clone signature for inversions. To ensure only split clones that signal a true inversion are detected, we also require read pair support for inversions (Medvedev *et al.*, 2009; Alkan *et al.*, 2011), and we discard any split clones that are not supported by read pairs. This step of the algorithm runs in *O*(*m* + *n*), where *m* is the number of read pairs with inversion signature and *n* is the number of split clones.

Each pair of split clones gives a hint about the existence of an inverted haplotype. There may be many incorrectly identified split clone inversion signatures, or a short clone may have multiple potential “mate”s with similar properties. Therefore, clustering multiple split clone pairs that share inversion breakpoint locations and inversion lengths can help resolve the inversion breakpoints more accurately (Figure 2c). To both resolve ambiguities from multiple possible split clone pairings, and unambiguously identify inversions, we construct an undirected graph, where each PSC is a node, and an edge between two nodes indicates that share predicted breakpoints.

We initially formulated the inversion detection using split clones as a Set-Cover problem similar to VariationHunter, however, we observed in both simulation and real data sets that due to segmental duplications and deletions around the breakpoints, Set-Cover approximation selected only one of the inversion breakpoints correctly (Supplementary Note). We therefore formulate the problem as finding maximal quasi cliques in the inversion cluster graph. This formulation allows existence of incomplete clusters, and tolerates some split clones to be included in a true cluster, and as a result, increases flexibility and avoids getting stuck in a local optimum.

We construct a graph *G* = (*V, E*) as follows. Each node in the graph denotes a PSC, as explained above, and each PSC will therefore represent a potential pair of inversion breakpoints. We put an edge between two nodes if the two representative PSCs agree with breakpoint locations through simple intersection (they are compatible with each other). Formally,

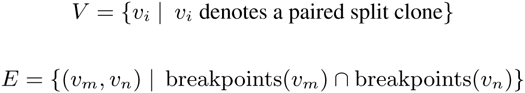

To find an approximate solution for the maximal quasi clique problem, we use an approximation algorithm previously suggested by Brunato *et al.* (2008), and we set the *tabu, γ*, and *λ* parameters to |graph|/10 *rounds*, 50%, and 60%, respectively. We obtained the values for these parameters by another grid optimization on experimental graphs depicting worst case scenarios (Supplementary Note).

When a quasi clique is found, the nodes within the clique denote a set of PSCs that are clustered together to mark an inversion. The breakpoint of this cluster is obtained by intersecting its split clones using a heuristic based on read pair support and the gap size. Next, the read pair support for the breakpoints within a distance is recalculated using the discordant read pairs. All clusters are then checked for any overlap on one side of the breakpoints and only the one with larger read support to split clone support ratio is kept and the rest are discarded. We propose to use this ratio to ensure fairness for less covered regions due to either random mapping or repeated regions. We report the final clusters after removing those that intersect with duplications and assembly gaps (>40%). A flowchart summarizing the dipSeq algorithm is available in the Supplementary Note.

### 4.5 Experimental validation

We tested the presence of an inversion in the cell line of the NA12878 individual predicted to carry an inverted haplotype. For this purpose, we used metaphase fluorescent in situ hybridization (FISH) validation for inversions larger than 2 Mbp using two probes located inside of the inversion. Similarly, we used interphase triple-color FISH to validate inversions smaller than 2 Mbp and larger than 500 Kbp using two probes inside and one outside the inversion.

## 5 RESULTS

We applied our algorithm to discover large inversions using three simulated and one real data set. The first simulation aims to both estimate the minimum read coverage requirements for accurate reconstruction of clones, and the effectiveness of our algorithms in large inversion discovery. We designed the second simulation to understand how dipSeq behaves in the presence of other structural variants that may have similar split clone signature. The third simulation depicts the robustness of dipSeq to segmental duplications. We finally used dipSeq to discover large inversions in the genome of an individual of Northern European descent (NA12878). For the real data, we compared our results with the InvFEST database of known inversions (Martínez-Fundichely *et al.*, 2014), and we applied experimental validation for the novel inversion calls.

### 5.1 Simulation Experiments

We designed three simulation experiments to test and demonstrate the power of dipSeq for inversion discovery. The details of each experiment and results are given in the Supplementary Note.

#### Simulation 1

In order to test the correctness of dipSeq, first, we randomly implanted 8 large inversions (500 Kbp to 10 Mbp) to the human reference genome (GRCh37) chromosome 1. Half of the simulated inversions were homozygous, and the remaining were heterozygous. We then randomly selected BAC-sized intervals (*μ* = 150 Kbp, *σ* = 40 Kbp) from both chromosome 1 homologs at ~3X physical coverage, which we randomly placed into 288 pools and simulated paired-end reads of length 100 bp (fragment size *μ* = 600 bp, *σ* = 60 bp) using `wgsim`^1^. We generated three different data sets at 3X, 5X, and 10X depth of coverage to investigate the effect of read depth on our inversion calls. We mapped the reads to the reference genome using both BWA and mrFAST aligners and applied our clone reconstruction method. We were able to correctly infer 87.18% and 86.40% of the clones that were not located on the breakpoints using the BWA and mrFAST alignments, respectively (Supplementary Note). Using the inferred clones, dipSeq could find all 8 inversions. It performed similarly in terms of sensitivity at all levels of depth of coverage, and returned no false positives.

#### Simulation 2

As a second simulation test, we explored dipSeq’s performance when there are other forms of structural variation close to or intersecting the inversion breakpoints, therefore emulating complex rearrangements. We used the same simulated inversions, and we additionally implanted deletions and duplications. We also inserted two additional inverted duplications to test whether dipSeq would predict them as normal inversions. We then repeated our clone and paired-end read simulation as explained above. However, due to random simulation, one of the inversion breakpoints was not “detectable” i.e. no clones spanned the breakpoint. After clone reconstruction, dipSeq was able to find all remaining 7 inversions correctly even at 3X sequence coverage. In addition, dipSeq did not incorrectly identify inverted duplications as bona fide inversions.

We further tested the efficacy of using whole genome sequencing (WGS) based inversion discovery algorithms on this data. For this purpose, we simulated WGS data sets, again using `wgsim`, at 3X, 5X, and 10X from the same chromosome homologs with the implanted inversions and SVs. We mapped the simulated reads to the reference human genome (GRCh37) with both BWA and mrFAST, to test the detection performance of three algorithms: INVY (Rausch *et al.*, 2012), LUMPY (Layer *et al.*, 2014), and VariationHunter (Hormozdiari *et al.*, 2009). We used the BWA alignments for INVY and LUMPY, and mrFAST alignments for VariationHunter, as per each tool’s usage recommendations. As expected, INVY and LUMPY failed to discover any of the implanted inversion events, as they are mainly designed for finding shorter inversions. VariationHunter was able to identify only one inversion out of 8, which may be due to VariationHunter’s ability to incorporate all map locations, and a higher maximum inversion size threshold.

#### Simulation 3

In the third experiment, we tested the robustness of dipSeq to segmental duplications, by implanting 4 large inversions (100 Kbp to 5 Mbp) to human chromosome 22, where the breakpoints intersect with segmental duplications. Two of the simulated inversions were homozygous, and the remaining were heterozygous. In addition, one of the inversions was placed near an assembly gap. We then randomly selected both BAC size (*μ* = 150 Kbp, *σ* = 40 Kbp) and fosmid size *(μ* = 40 Kbp, *σ* = 10 Kbp) intervals from both chromosome 22 homologs at ~4X physical coverage, which we then randomly placed into 288 pools ensuring that the clones do not span the unmapped areas.

We then simulated paired-end reads of length 100 bp (fragment size *μ* = 600 bp, *σ* = 60 bp) using `wgsim` and generated three different data sets at 3X, 5X, and 10X depth of coverage, for both BAC and fosmid simulations. Next, we mapped the reads to the entire reference genome using the BWA aligner, and finally applied dipSeq. Our algorithm was able to precisely detect all four inversions in each experiment, and returned no false positive predictions. We noticed that increasing the sequence coverage did not improve the results, however, when the physical coverage was reduced to 3X, some inversions became undetectable since no clones spanned their breakpoints.

### 5.2 Real data set from NA12878

Next, we tested dipSeq using a real pooled clone sequencing data set generated from the genome of NA12878. We mapped paired-end reads from a total of 288 pools (Methods) using both BWA and mrFAST to the reference genome. Average fragment length of the paired-end reads was ~450 bp, with a standard deviation of ~98 bp. Using our algorithms, we reconstructed the clone locations, which showed an average clone length of ~140 Kbp and a standard deviation of ~40 Kbp.

For inversion discovery, we set the minimum and maximum inversion size thresholds as 500 Kbp and 10 Mbp, respectively. Although it is theoretically possible to detect inversions as small as a typical clone size (150-200 Kbp), we chose the minimum size as 500 Kbp due to the limitations of the FISH method we used for validation (Methods). After the initial split clone clustering and maximal quasi clique approximation (Methods), we filtered those inversion clusters without read pair signature support. We generated two main callsets using BWA and mrFAST, where >83% of the calls were shared. We then randomly selected a total of 11 inversions for experimental validation (Table 1). We then compared our predictions with the known inversions reported in the InvFEST database (Martínez-Fundichely *et al.*, 2014), and found that dipSeq could correctly identify all three inversions that are previously *validated* in the genome of the same individual; a 5 Mbp inversion in 8p23.1 (Antonacci *et al.*, 2009), a 1.5 Mbp inversion in 17q12 (Antonacci *et al.*, 2009), and a 2 Mbp inversion in 15q13.3 (Antonacci *et al.*, 2014) (Table 1). Out of the remaining 8 inversion predictions, 2 could not be tested due to the segmental duplications around the breakpoints. We tested the remaining using FISH experiments (Methods), and validated a novel inversion in the 15q25 locus (Figure 3a,b). We also show the visualization of a previously characterized 15q13.3 inversion (InvFEST ID: HsInv 1049) using the SAVANT browser (Fiume *et al.*, 2012) in Figure 3c. We used dipSeq with different parameters and generated two more data sets, which are not extensively tested (Supplementary Section 1.14).

**Fig. 3.**
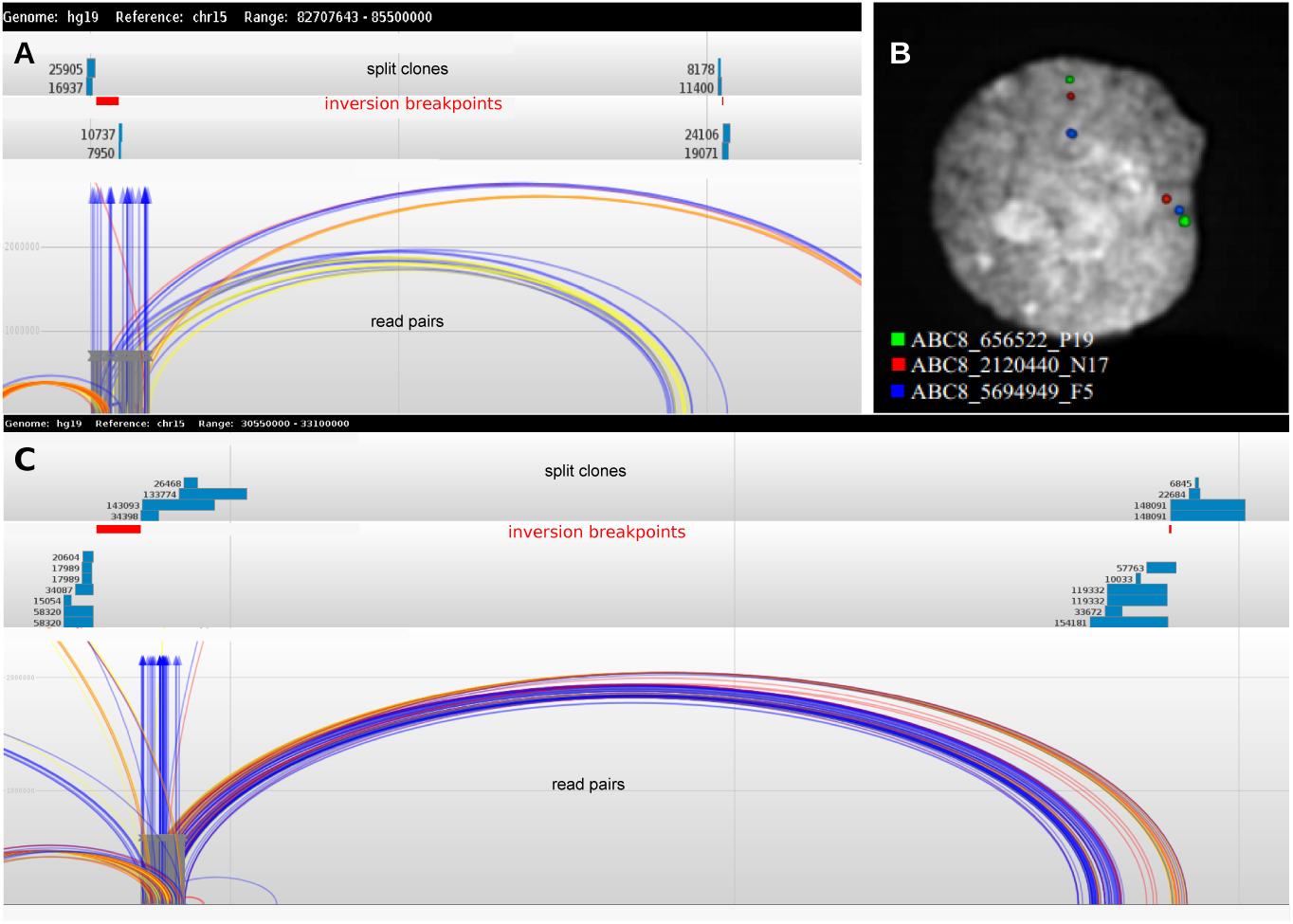
Inversions discovered by dipSeq in the NA12878 genome. (A) Novel inversion found at chr15:83,089,659-84,865,500 (inner coordinates). We show the locations of split clones and the supporting read pairs using the SAVANT browser (Fiume *et al.*, 2012). (B) Experimental validation of the novel inversion discovered using interphase FISH (green-red-blue: direct, green-blue-red:inverted). (C) SAVANT browser view of the previously known inversion at chr15:30,433,406-32,898,559. SAVANT read pair colors are as follows. Light blue: concordant, red: discordant by length, dark blue: one end inverted, yellow: everted (tandem duplication), gray: one end unmapped.

**Table 1.**
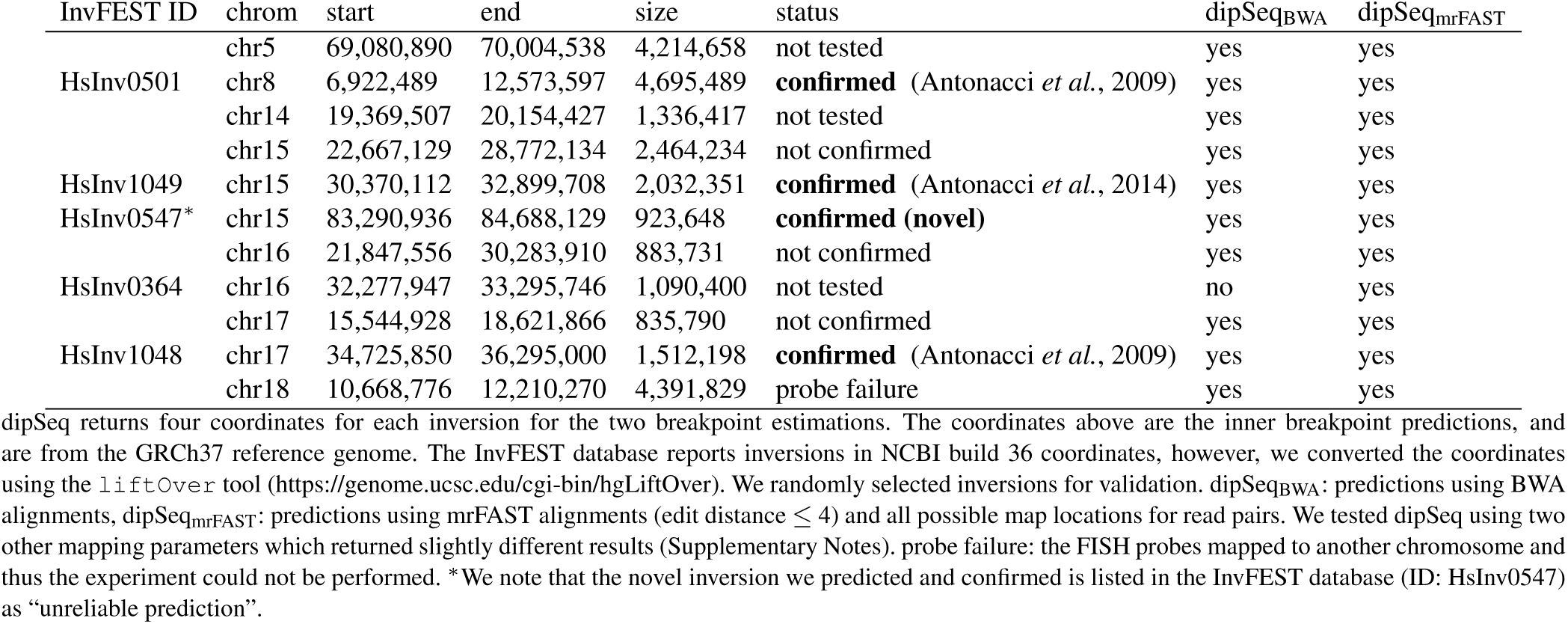
Summary of validation of inversions predicted in the genome of NA12878 using dipSeq.

## 6 DISCUSSION

In this paper, we presented a novel algorithm, dipSeq, to characterize large genomic inversions using a new sequencing method initially developed to improve haplotype phasing. Although it suffers from high false positive rate using real data (Table 1), dipSeq was able to identify all previously validated inversion events, and also discover a novel variant. Furthermore, dipSeq performed better with simulated data, suggesting that the relatively poor performance with the NA12878 genome may be improved with higher depth of coverage.

There are multiple directions that we can take to further improve dipSeq. First, to reduce the false discovery rate, we can incorporate split read sequence signature (Ye *et al.*, 2009), and we can perform local *de novo* assembly around the predicted breakpoint intervals with an approach similar to TIGRA (Chen *et al.*, 2014). However, since both of these methods need high sequence coverage, they might not be suitable to directly apply to the low-coverage data set we used. Instead, it will be better to simultaneously use WGS data generated from the genome of the same individual. Since the PCS method also requires WGS data for haplotype phasing, it can be expected to generate matching PCS-WGS data sets from the same genomes.

Another future research on dipSeq will be testing and improving its abilities to discover smaller, yet still large inversions (>100 Kbp). In this paper, we focused on inversions larger than 500 Kbp, because the upper size limit for GASVPro (Sindi *et al.*, 2012) algorithm is 500 Kbp, and only such large inversions can be reliably tested using FISH. Note that validating smaller inversions is a more difficult task, using fiber FISH, or PCR if the breakpoints lie within unique regions. In addition, the clone size distribution should be tighter to ensure clone reconstruction method does not artificially “merge” split clones into a single interval. Alternatively, we can try to use smaller clones such as fosmids, despite their limitations. We still would like to investigate dipSeq’s performance using real fosmid data, however, this may require additional algorithmic enhancements especially in the presence of nearby segmental duplications (Kitzman *et al.*, 2011). In this paper we present fosmid simulation experiments, and there is currently only one pooled fosmid sequencing dataset (Kitzman *et al.*, 2011) generated from the genome of a Gujarati Indian individual (NA20847). We would like to apply dipSeq to the NA20847 dataset and evaluate its performance with experimental validation.

dipSeq can also be extended to characterize other forms of large structural variation, including deletions, insertions, direct and inverted duplications. Each of these types of SV present themselves with different split clone signatures that we summarize in Supplementary Figure 6. We also note that, determining the location of a segmental duplication event is yet a largely unsolved problem, even when long reads are used (Chaisson *et al.*, 2015). It may also be possible to discover translocations using split clones, however, chance of finding incorrect split clones will also increase, causing a reduction in the performance of maximal quasi clique approximation.

In summary, dipSeq is the first algorithm that can discover large genomic inversions using high throughput sequencing technologies. Our understanding of the phenotypic effects of inversions is still limited, and one of the reasons of this is the lack of reliable and cost effective methods to characterize such events. This is also true for other complex rearrangements such as duplications and translocations. Improvements in characterization of large complex rearrangements will help us better understand the biological mechanisms that lead to phenotypic difference, disease, and evolution.

## AUTHOR CONTRIBUTIONS

C.A., F.A., and E.E.E. designed the study. C.A. and M.E.R developed the dipSeq algorithm. M.E.R. implemented and applied dipSeq to simulation experiments and real data. J.T. and C.T.A. built BAC clones, M.V. and F.A. generated pooled clone sequencing data. G.C. and M.M. performed validation experiments. C.A., F.A., and M.E.R. wrote the paper.

## ACKNOWLEDGMENTS

We would like to thank J. Kitzman, and B. Dumont for data access and their valuable input for the clone reconstruction algorithm, and J. Huddleston for computational assistance. We also thank M. C. Orhan for the discussions on probability calculations for clone reconstruction, B. Genç and M. Sarı for their help with the initial formulations of the problem, C. Arbib for discussions on how to formulate inversion events with estimated breakpoint intervals with Set-Cover, and M. Cáceres and S. Casillas for their help with the InvFEST database.

### Funding

Funding for this project was provided by a Marie Curie Career Integration Grant (303772) and an EMBO grant (IG-2521) to C.A., an NIH grant (HG004120) to E.E.E., and a Firb-Programma “Futuro in Ricerca” grant (RBFR103CE3) to M.V. C.A. also acknowledges support from The Science Academy of Turkey, under the BAGEP program.

### Conflict of Interest

E.E.E. is on the scientific advisory board for <u>DNAnexus, Inc.</u>

## Footnotes

1 https://github.com/lh3/wgsim

